# Lessons in genome engineering: opportunities, tools and pitfalls

**DOI:** 10.1101/710871

**Authors:** Ingrid Poernbacher, Sam Crossman, Joachim Kurth, Hisashi Nojima, Alberto Baena-Lopez, Cyrille Alexandre, Jean-Paul Vincent

## Abstract

CRISPR/Cas technology allows the creation of double strand breaks and hence loss of function mutations at any location in the genome. This technology is now routine for many organisms and cell lines. Here we describe how CRISPR/Cas can be combined with other DNA manipulation techniques (e.g. homology-based repair, site-specific integration and Cre or FLP-mediated recombination) to create sophisticated tools to measure and manipulate gene activity. In one class of applications, a single site-specific insertion generates a transcriptional reporter, a loss-of function allele, and a tagged allele. In a second class of modifications, essential sequences are deleted and replaced with an integrase site, which serves as a platform for the creation of custom reporters, transcriptional drivers, conditional alleles and regulatory mutations. We describe how these tools and protocols can be implemented easily and efficiently. Importantly, we also highlight unanticipated failures, which serve as cautionary tales, and suggest mitigating measures. Our tools are designed for use in *Drosophila* but the lessons we draw are likely to be widely relevant.

**AUTHOR SUMMARY:** The genome contains all the information that an organism needs to develop and function throughout its life. One of the goal of genetics is to decipher the role of all the genes (typically several thousands for an animal) present in the genome. One approach is to delete each gene and assay the consequences. Deletion of individual genes is now readily achieved with a technique called CRISPR/Cas9. However, simple genetic deletion provides limited information. Here we describe strains and DNA vectors that streamline the generation of more sophisticated genetic tools. We describe general means of creating alleles (genetic variants) that enable gene activity to be measured and experimentally modulated in space and time. Although the tools we describe are universally applicable, each gene requires special consideration. Based on our experience of successes and failures, we suggest measures to maximise the chances that engineered alleles serve their intended purpose. Although our methods are designed for use in Drosophila, they could be adapted to any organism that is amenable to CRISPR/Cas9 genome modification.

## INTRODUCTION

During the past century, biologists have largely relied on forward genetics to determine gene function. This involved identifying, from a large number of randomly generated mutations, those that affect a particular phenotype. This has been a powerful gene discovery tool. For example, forward genetics has identified most of the genes involved in segmental patterning of *Drosophila* ([1]) and many have turned out to be broadly conserved and hence of general interest. Metazoan reverse genetics (from gene to phenotype) was initiated in mice with gene targeting by homologous recombination ([2]), allowing the role of any gene to be assessed albeit in a laborious way. Genome-wide reverse genetics then became possible, particularly in worms and flies, with the development of RNAi technology ([3]). Despite its ease, RNAi has limitations, such as incomplete knockdown. True genetic null alleles remain the gold standard to determine whether a gene is essential for a particular function. Thanks to CRISPR/Cas technology, generating such mutants has become within the reach of any research laboratory. Indels are readily created by inducing two site-specific double strand breaks (DSBs), thus deleting the intervening genomic sequences ([4]). With this approach, any gene can be inactivated in most model organisms and the phenotypic consequences assessed.

Although undoubtedly powerful, null alleles are limited. For example, they do not readily allow phenotypic analysis of essential genes beyond their lethal phase. In mice, this limitation is overcome by conditional alleles, whereby an essential exon is excised by Cre-mediated recombination ([5]). In *Drosophila*, delayed gene inactivation is traditionally achieved in clones generated by mitotic recombination ([6]). Indeed, clonal analysis has led to major insight over the years. However, it does not readily lend itself to making whole mutant tissue or organ without the possible complication of cell-lethal sister clones. Another shortcoming of classic clonal analysis is that it is not suitable to remove gene activity from post-mitotic cells or for genes that are located near centromeres. As described in this paper, CRISPR/Cas technology not only allows the creation of conditional alleles but also opens the way to the generation of alleles that enable gene activity to be measured and manipulated in a wide variety of ways. We first describe protocols and vectors to generate authentic transcriptional reporters as well as tagged gene product expressed from the endogenous locus. We then provide guidance for making insertion-ready deletion alleles (deletion of essential sequence and replacement with an attP integrase site) and argue that such alleles constitute a multi-purpose platform for the subsequent creation of diverse alleles that report on gene function or enable gene manipulations with unparalleled sophistication. To be effective, these insertion-ready deletion alleles must fully inactivate gene function and at the same time allow native expression of reintroduced genetic elements. We describe instances when these aims unexpectedly failed and provide guidelines on how to avoid such mistakes. Although our protocols are designed for *Drosophila*, most of the lessons we draw from failure apply generally to the generation of sophisticated alleles in other organisms.

## RESULTS AND DISCUSSION

### Direct tag insertion

Site-specific insertion of exogenous DNA into the *Drosophila* genome has become routine thanks to CRISPR/Cas technology ([7, 8]). Typically, DSBs are induced and repaired by exogenously supplied DNA comprising homology arms flanking the fragment to be inserted (the repair fragment). This is practically achieved in *Drosophila* by co-injecting the repair fragment and a plasmid expressing gRNAs into the germ line of a Cas9-expressing strain. Many such strains are available but they tend to be relatively unhealthy. We recently found that, fortuitously, expression of Cas9 fused to a monomeric streptavidin (Cas9-mSA), which was recently shown to be effective in mice ([9]), is healthier than the original *nos-Cas9* stocks, while still being effective at inducing DSBs. We therefore routinely use *nos-Cas9-mSA* transgenics (See Table 1) as our host strain of choice. For the introduction of small epitope tags, the repair fragment can be commercially synthesized as a singlestranded biotinylated oligonucleotide (with homology arms of about 60 bases flanking DNA encoding the tag; [10]). Although this method requires little molecular biology at the outset, it involves significant effort for PCR screening of candidate integrants. Recognizable genetic markers obviate the need for PCR screening but, because of their size, are not compatible with synthetic repair DNA. Instead, DNA encoding visible genetic markers needs to be incorporated in a gene-targeting plasmid. We have therefore created targeting vectors that use a Cre-excisable *Pax-Cherry* cassette (strong Cherry expression in the eyes) as a genetic marker (Fig. 1 and Table 1). As described below, these plasmids, enable the insertion of 2xHA (small tag), GFP, or Scarlet at specific locations in the genome.

**Fig. 1.**
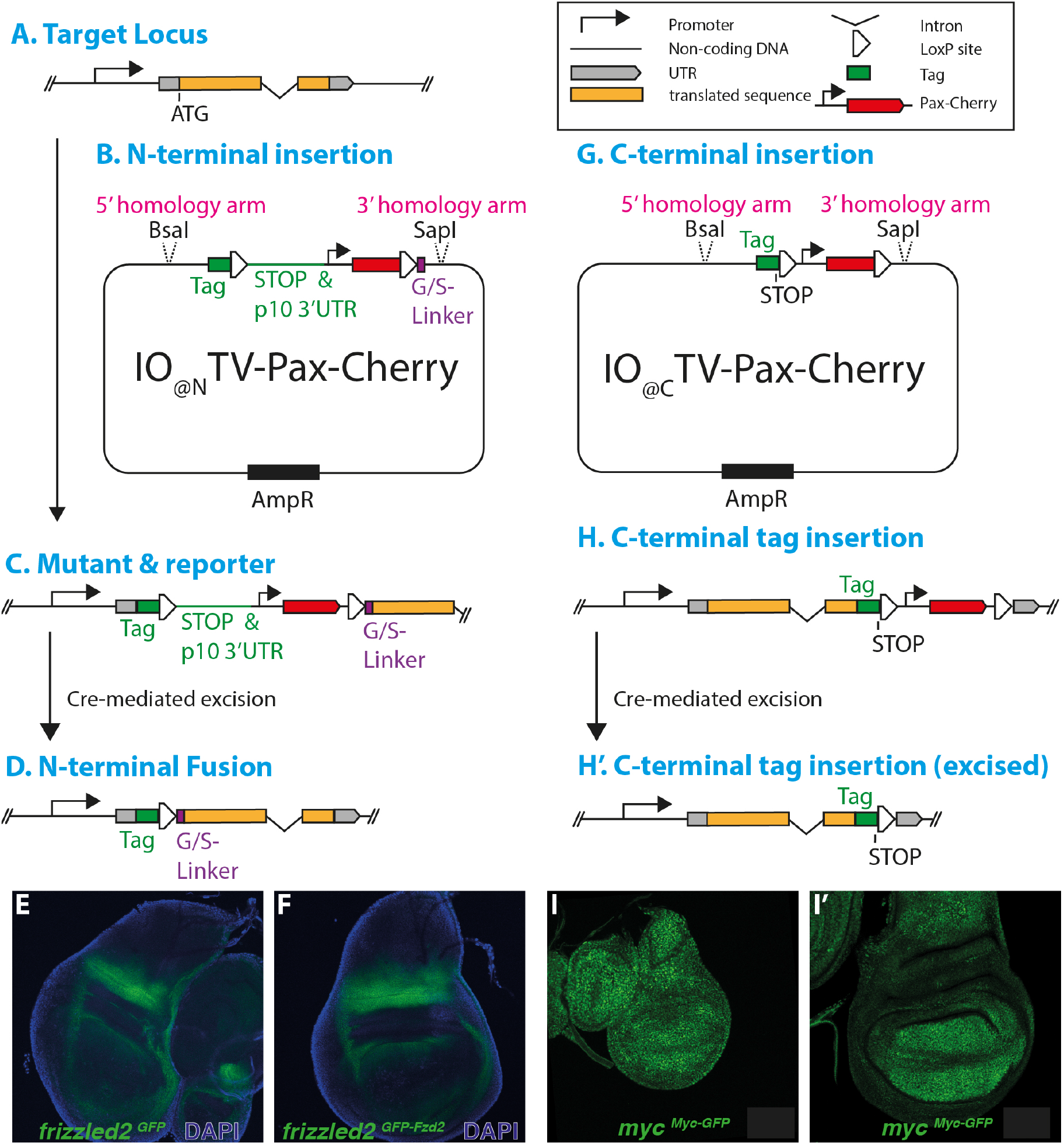
Creation of insertion-only alleles. (A) Diagram of a sample locus, with coding exons in orange and untranslated sequences in grey. (B) Targeting vector for N-terminal fusion (Insertion Only; IO_GFP@N_TV-Pax-Cherry; IO_2xHA@N_TV-Pax-Cherry or IO_Scarlet@N_TV-Pax-Cherry) highlighting key features. Green box indicates the tag (2xHA, GFP or Scarlet). This is followed by a LoxP, a termination codon and p10 3’UTR (not for 2xHA). White triangles indicate LoxP sites. Upon Cre-mediated excision, the tag is brought in frame with the gene’s coding region with an intervening linker (G/S linker) and residual LoxP site. Red box indicates Cherry coding region of *Pax-Cherry*. Unique restriction sites for insertion of the homology arms are also shown. Following insertion of suitable homology arms, this vector is co-injected in a Cas9-expressing strain, with a vector expressing gRNAs that target the translation start (e.g. pCFD3 or pCFD4). (C) Following integration, the resulting allele harbours the Cre-excisable region comprising the termination codon, the p10 3’UTR and the *Pax-Cherry* cassette. This is expected to disrupt expression of the targeted gene. At the same time, the reporter (GFP or Scarlet) is expressed in the pattern of the targeted gene (a 3’UTR was not included downstream of 2xHA since IO_2xHA@N_TV-Cherry is not intended to be used for the generation of a transcriptional reporter). (D) Cre-mediated excision fuses the reporter to the 5’ end of the coding sequence, leading to expression of a N-terminal fusion protein with an intervening LoxP and a stretch of 10 Gly-Ser (G/S linker). (E) Expression of GFP in a wing imaginal disc at third instar stage following insertion of an integration-only vector in *frizzled2* (*fzd2*) (just downstream of the sequences encoding the signal peptide). This results in expression of secreted GFP in the *frizzled2* pattern. Note that this *frizzled2^GFP^* transcriptional reporter was generated with the general strategy outlined in (A-C) but with an early version of the IO_GFP@N_TV-Pax-Cherry vector lacking the p10 3’UTR. (F) GFP expression from the same allele after Cre-mediated conversion. Now a GFP-Frizzled2 fusion protein is expressed. (G) Targeting vector for C-terminal fusion of GFP or 2xHA (IO_GFP@C_TV-Pax-Cherry or IO_2xHA@C_TV-Pax-Cherry). Here the tag is followed by a termination codon and *Pax-Cherry* as a genetic marker. Only *Pax-Cherry* is flanked by LoxP sites. As in B, the targeting vector is co-injected with pCFD3 or pCFD4 expressing a gRNA that targets the STOP codon. (H, H’) Diagram of locus after integration of a IO_2xHA@C_TV-Pax-Cherry construct before (H) and after Cre-mediated excision (H’). (I, I’) Myc-GFP fusion expression from an allele generated as in G-H in a wing imaginal disc at second (I) and third (I’) instar.

**Table 1.** Tools

|  |  |  |
| --- | --- | --- |
| *nos-Cas9-mSA* | Healthy *Drosophila* host strain for injection. Available with landing sites on 2d (attP40) and 3d chromosome (attP2). Carries Pax-GFP as a genetic marker. | This study |
| CFD4 & CFD4 | Vectors for expression of gRNA | [18] |
| IO@N<sub>TV</sub>-Pax-Cherry (Targeting Vector for Insertion-Only at N-terminus). Variants: IO<sub>GFP</sub>@N<sub>TV</sub>-Pax-Cherry IO<sub>Scarlet</sub>@N<sub>TV</sub>-Pax-Cherry IO<sub>2xHA</sub>@N<sub>TV</sub>-Pax-Cherry | To generate LOF mutant, transcription reporter, and (after Cre-mediated recombination) N-terminal fusion protein. Variants for GFP (with p10 3'UTR), Scarlet (with p10 3'UTR), and 2xHA (without p10 3'UTR). Genetic marker = Pax-Cherry. | This study (see [19] for parent plasmid) Fig. 1 |
| IO@c<sub>TV</sub>-Pax-Cherry (Targeting Vector for Insertion at C-terminus) Variants: IO<sub>GFP</sub>@c<sub>TV</sub>-Pax-Cherry IO<sub>2xHA</sub>@c<sub>TV</sub>-Pax-Cherry | To generate C-terminal fusion protein. Variants for GFP and 2xHA. Genetic marker = Pax-Cherry. | This study (see [19] for parent plasmid) Fig. 1 |
| TV<sub>ΔattP</sub>-Pax-Cherry (attP deletion targeting vector) | Targeting vector to generate insertion-ready deletion alleles. Genetic marker = Pax-Cherry. | This study Fig. 2 |
| RIV<sub>attB</sub>-Pax-GFP (Generic re-integration vector) | To re-integrate genetic elements at attP site of insertion-ready deletion alleles. Genetic marker = Pax-GFP.<br>This complements RIV<sub>attB</sub>-Pax-Cherry and RIV<sub>attB</sub>-white+ described in [13] | This study Fig. 3 |
| RIV<sub>FRT.attB.FRT</sub>-Pax-GFP (Re-integration vector for FRT-flanked fragments) | To re-integrate FRT-flanked genetic elements at attP site of insertion-ready deletion alleles. Genetic marker = Pax-GFP.<br>This complements RIV<sub>FRT.attB.FRT</sub>-Pax-Cherry and RIV<sub>attB</sub>-white+ described in [13] | This study Fig. 3 |

We first describe plasmids for insertion of DNA encoding GFP or Scarlet at the 5’ end of the gene of interest (Fig. 1A-D). Typically, the CRISPR site and homology arms are chosen so that the first codon of GFP (or Scarlet) corresponds to the translation start of the endogenous gene, thus enabling expression under authentic control of the endogenous gene. The 3’UTR of p10 ([11] [12]) was included to boost RNA stability, thus maximizing expression. Following integration, the p10 3’UTR, as well as the transcriptional termination signal of the *Pax-Cherry* cassette, interrupt normal transcription and translation of the endogenous gene. Therefore, integration of this targeting construct is expected to inactivate gene function (however see pitfalls in the next paragraph), while at the same time creating a reporter gene. A *frizzled2^GFP^* reporter created by this general strategy is shown in Fig. 1E. Such alleles can be readily converted to a tagged protein reporter by excision of the *Pax-Cherry* cassette (and the p10 3’UTR) with a Cre-expressing transgene. In the resulting strain, the endogenous promoter drives expression of the fluorescent protein fused to the N-terminus of the protein of interest, with intervening amino acids encoded by the residual LoxP site and an additional G/S linker (i.e. GFP-Frizzled2 fusion protein in Fig. 1F). We found no adverse effect of this “LoxP linker” although it is obviously essential that the homology arms are chosen to ensure that the tag is in frame with the gene’s initiation codon. A similar plasmid was constructed for insertion of 2xHA but, in this case, the p10 3’UTR was omitted because 2xHA would only be used as a tag (after removal of the *Pax-Cherry* cassette) and not as a transcriptional reporter. Note that for most secreted proteins (such as Frizzled2), the tag must evidently be inserted downstream of the signal peptide (see http://phobius.sbc.su.se), not at the translation initiation codon, to ensure that the fusion protein traffics through the secretory pathway. One could also envision inserting the tag further downstream to create an internal fusion although we have not yet attempted such a modification.

The above procedure suffers from a potential pitfall that needs to be mentioned. In theory, our N-terminus-targeted insertion-only targeting vector can generate, from a single integrant, a transcriptional reporter, a loss of function mutant, and a protein reporter. However, it is important to note that the mutant may not always be null. This is because one cannot exclude the possibility of transcription/translation reinitiation downstream of the *Pax-Cherry* cassette. Although unlikely, this scenario must be kept in mind during phenotypic analysis. In doubt, it might be safer to delete additional genomic sequences, as described in the subsequent section.

To complement the above vectors, we have also created plasmids for direct insertion of a tag at the 3’ end of the coding region (Fig. 1G, H). Here GFP or 2xHA is inserted at, or shortly upstream of, the endogenous stop codon. As before, selection of successful integrants is achieved by the presence of a downstream excisable *Pax-Cherry* cassette. In this case, the cassette does not interrupt the endogenous reading frame. However, it does separate the coding region from the endogenous 3’UTR and this could have an impact on expression levels. Excision of the *Pax-Cherry* cassette will restore the position of the endogenous 3’UTR (albeit with an intervening residual LoxP site), ensuring near native expression of the tagged protein as illustrated for *Myc-GFP* (Fig. 1I).

Insertion-only alleles are so easily recovered that this approach can be extended to generating reporter genes from a transgenic BAC. This is particularly useful for genes that are haplo-insufficient or to generate reporters at a different chromosomal location from that of the endogenous gene. The strategy we suggest is to first generate transgenic flies carrying a BAC containing all essential regulatory elements (this can be confirmed by rescue of a pre-existing null mutant). Once in the genome, the transgene can be readily modified with the insertion-only targeting vector described above, bearing in mind that the endogenous locus will also be targeted.

The genetic manipulations described above allow rapid insertion of tags/reporters in any locus. One key advantage of insertion-only modifications (no deletion) is that they occur at high efficiency by comparison to insertions combined with deletion (especially for large deletions, e.g. over 2kb). Another significant advantage is that one event generates multiple tools. This could be particularly useful for species that have a longer generation time or are less amenable to targeted homologous recombination than *Drosophila melanogaster*. However, one possible shortcoming is that insertion-only modifications do not always completely remove gene activity. Below, we describe insertion-ready deletion alleles that are more likely to abrogate gene activity while enabling a wide range of genetic manipulations.

### Design of insertion-ready deletion alleles (ΔattP)

Insertion-only alleles can rapidly provide insight about gene expression and function. However, more extensive genetic analysis requires a two-step approach with the generation, as a first step, of a deletion with concomitant insertion of an integrase site. This can be achieved with the targeting vector illustrated in Fig. 2 B, which also allows the mutation to be tracked with *Pax-Cherry*. The vector contains two multiple cloning sites for insertion of homology arms. Here, it is worth mentioning that inclusion of the CRISPR target site at the distal end of one homology arm improves targeting efficiency by ensuring linearization of the targeting vector. The insertion-ready deletion alleles constitute a foundation on which a variety of genetic tools can be built since they are amenable to subsequent insertion of various genetic elements such as transcriptional reporters, epitope tagged cDNA, rescuing constructs, etc. (see also [13]). Although this approach is powerful and versatile, it is liable to unanticipated problems, which can be avoided with careful design of the original deletion. Two constraints must be considered in designing insertion-ready deletion (ΔattP) alleles: the need for the deletion to completely inactivate gene activity and the requirement that DNA fragments integrated in the attP site be transcribed in the same pattern as the endogenous gene. In practice the latter is ensured by judicious choice of the 5’ breakpoint, typically upstream of the translation initiation codon, while the former is determined by the 3’ breakpoint, i.e. the size of the deletion. We address these two constraints in turn, starting with the size of the deletion required to abrogate gene activity.

**Fig. 2.**
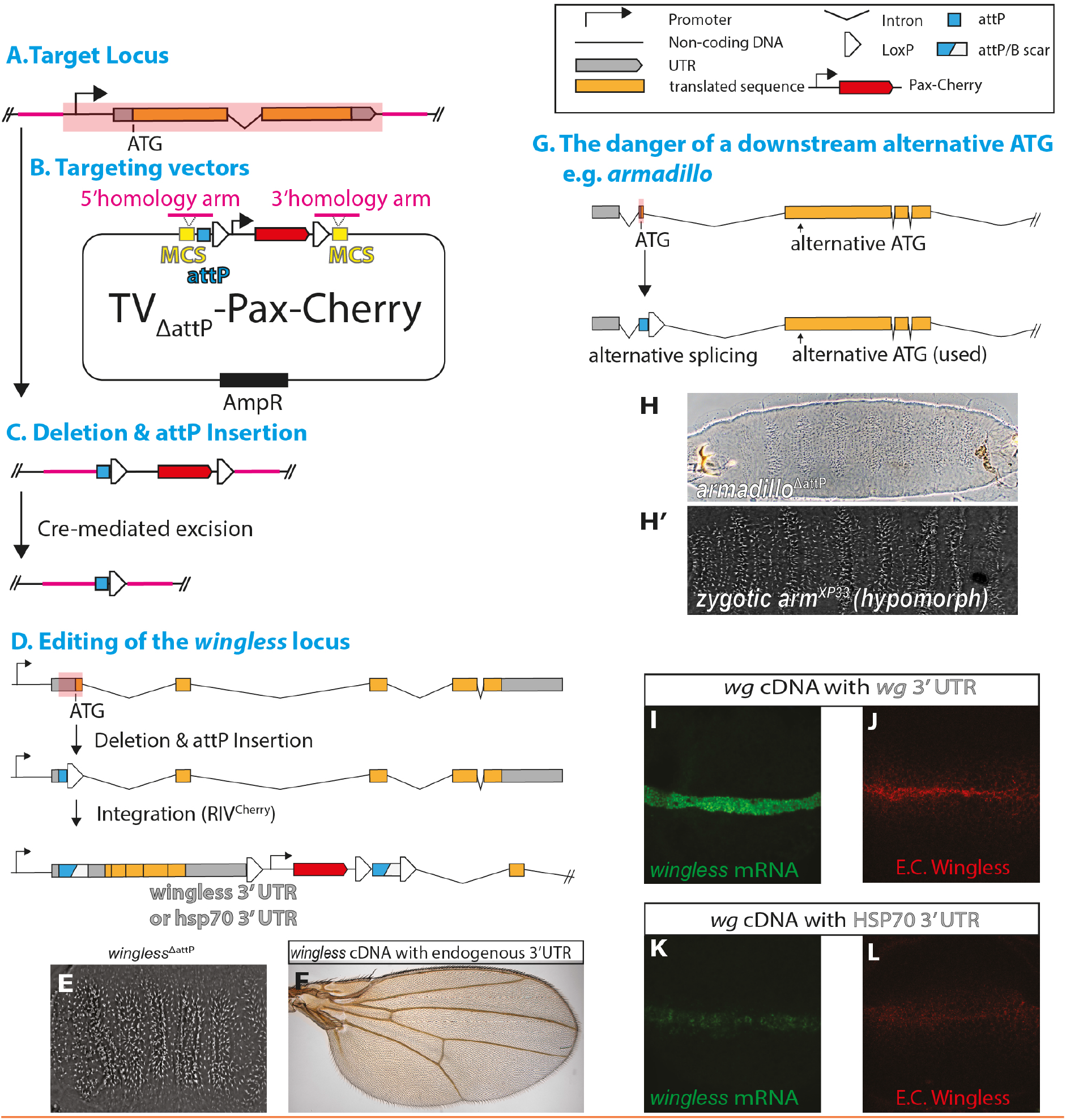
Creation of insertion-ready deletion alleles (ΔattP) (A) Diagram of a generic locus to be modified, with coding exons in orange and untranslated sequences in grey. Location of homology arms are indicated in magenta, with the intended deletion marked with a red box. In this example the deletion spans the entire target gene, including the 5’and 3’UTRs, and extends upstream of the TATA box (into an area of low conservation). This strategy will ensure complete loss of function and avoids any disturbance of translation initiation, thus maximising the chance of faithful expression of re-integrated material. Such an approach is therefore the safest but not always achievable for large genes. (B) Diagram of the targeting vector, TV_ΔattP_-Pax-Cherry, which is an improved and decluttered version of our previously published pTV^Cherry^ ([13]). Key features include the attP site, which will allow phiC31 integrase-driven re-integration, and an excisable *Pax-Cherry* cassette flanked by LoxP sites. Genomic deletions are achieved by co-injecting Δ_attP_ TV-Cherry with a vector expressing the two guide RNAs targeting the deletion breakpoints (pCFD4). (C) Diagram of the resulting allele, showing the genomic deletion and the inserted *Pax-Cherry* cassette, which can be removed by a Cre-mediated excision. (D) Top: Diagram of the *wingless* locus, with coding exons in orange, untranslated sequences in grey and the deleted region marked with a red box. Middle: The resulting Δ_attP_ allele after Cre-mediated excision of the *Pax-Cherry* cassette. Bottom: The *wingless*^ΔattP^ locus after phiC31 integrase-mediated re-integration of a *wingless* cDNA (two different 3’UTRs were used, see below). Here re-integration was achieved with the *Pax-Cherry-marked* RIV^Cherry^ described in [13]. (E) Cuticle phenotype of homozygous embryos carrying the modification shown in the middle of D (deletion of the first coding exon). This is the same as that of null alleles ([20]). Re-integration of *wingless* cDNA with either the endogenous wingless 3’UTR or a hsp70 3’UTR (as shown at the bottom of D) rescues the *wingless* mutant phenotype as evidenced by the wild-type appearance of wings (F, data not shown). (G) Top: The *armadillo* locus with coding exons in orange, untranslated sequences in grey and the region deleted by homologous recombination marked by a red box. Bottom: The *armadillo*^ΔattP^ allele after Cre-mediated excision of *Pax-Cherry*. (H, H’) The deletion strategy outlined in (G) leads to partial loss of function as evidenced by cuticle phenotype (H) resembling that of a known hypomorphic allele (*arm^XP33^*, H’) ([21]). Residual activity is likely due to expression of truncated protein resulting from alternative splicing (G). (I-L) Expression levels of the re-integrated *wingless* cDNA are influenced by the nature of the 3’UTR. With the *hsp70* 3’UTR, expression (RNA and protein) is markedly reduced compared to expression with the native 3’UTR.

Since regions of up to 30kb can be deleted and replaced by an attP site (unpublished observation), the safest strategy for ensuring complete loss of function would be to delete the entire target gene (Fig. 2A-C). However, we find that, in practice, it is preferable to limit deletion size as much as possible. Smaller deletions (< 2 kb) are more efficiently obtained and less prone to unexpected rearrangements. In addition, small deletions are more amenable to rescue by re-integration of the deleted fragment since large deletions would require the reinsertion of long DNA fragments (to ensure restoration of all regulatory elements), which are cumbersome to manipulate. Nevertheless, it is necessary that the deletion be large enough to completely abrogate gene function. Whether this can be achieved with a relatively small deletion must be assessed on a case-by-case basis. For a protein earmarked to the secretory pathway, a small deletion will suffice since it is only necessary to remove the signal peptide-encoding exon to fully inactivates the gene (e.g. *wingless*, [14]; Fig. 2D, E). For other genes, deletion of the exon bearing the translation initiation site may not suffice since downstream translation initiation sites could become available. As a real-life example, we found that deletion of *armadillo*’s second exon, which contains the initiation codon, only caused partial loss of function, most probably because of *de novo* splicing of the first and third exons (Fig. 2G, H). As a rule, it is advisable to consider deleting all the in-frame translation initiation sites that are likely to produce a functional or partially functional protein. This may be difficult when such sites are located downstream of a large intron (since a large deletion would be needed). In such situations, it is worth considering to remove only one (or more) downstream essential exons, such as for example exons encoding the catalytic domains of enzymes. This strategy could ensure complete loss of function but would reduce the range of tools that can be generated by re-integration (for example the N-terminal portion left in the genome could not be modified). In summary, although complete abrogation of gene activity is often possible, this cannot be guaranteed for all genes. At times, compromise will be needed.

One important feature of insertion-ready deletion alleles is that they enable expression of reporters or modified forms of the coding region in a pattern and level that mimic those of the endogenous gene. If the mutation causes a well-defined phenotype, one way to ensure that re-integrated DNA fragments are correctly expressed is to test whether full gene activity is rescued by insertion of the deleted DNA or a cDNA into the attP site. For genes that have no obvious phenotype, this is not possible and the next best option is to verify that a reporter integrated in the attP site is expressed like the native gene. Note that, as described below, the breakpoints of the deletion must be carefully chosen to maximize the chance that re-integrated fragments are faithfully expressed (see section on re-integration of genetic elements for further details). For most essential genes we found that gene activity can be restored by re-integration of the deleted genomic DNA or in some cases, a cDNA (e.g. *wingless*, Fig. 2D, F). In a few instances however, rescue was not achieved. We suggest that such failures could be explained by interference with the 5’ UTR, the 3’UTR, a regulatory intron, or splicing, as described in further detail below.

The importance of preserving the native 3’UTR is illustrated by re-integration into the *wingless* locus. Here the gene was knocked out by deletion of the first exon, which harbors the signal peptide. This could be rescued by insertion of a *wingless* cDNA into the attP site ([14]), showing that all regulatory elements (including those located in downstream introns) were functional. We found however that, in this instance, it was important to bring back the native 3’UTR along with the cDNA for full rescue to take place. As shown in Fig. 2I-L, much weaker expression was seen with another 3’UTR (that of the *hsp70* gene). We suggest that the initial deletion should be designed in such a way that, upon re-integration, the native 3’UTR is used by the rescued fragment. Alternatively, the 3’UTR should be included in the rescue construct.

Most of the time, re-integration of the deleted fragment leads to restoration of gene function and expression. However, the small scar created by integration of an attB sequence into an attP site (referred to as attP/B) could have deleterious effects. Indeed, we encountered several instances when DNA fragments inserted into an attP site located just upstream of the translation initiation site failed to be expressed, even though they were located downstream of the transcription start site and were thus expected to be transcribed. For example, we found that a cDNA encoding HA-tagged Notum failed to be expressed (and to rescue the mutant phenotype) when inserted into a *notum*^ΔattP^ allele with the attP site located 8 nucleotides upstream of the ATG (unpublished observation). We suggest that the attP/B sequence could interfere with ribosome entry, transcription, and/or RNA stability, especially if it is located within 30 bp of the translation initiation codon. One solution is to make sure that the attP site (the deletion’s 5’ break point) is located further upstream (ideally more than 50 bp away from ATG) but still downstream of the transcription start site and in a region that is poorly conserved in other *Drosophila* species (as assessed at https://genome.ucsc.edu). In some cases, this may not be possible and it might be necessary to choose a 5’ breakpoint upstream of the transcription start site to find a suitable unconserved region (Fig. 2A). This can be a safer option although it requires reconstitution of the transcription start site and 5’UTR in the re-integrated fragments. Deletion of the whole 5’UTR may also increase the chance that the deletion fully abrogates gene activity. However, the price to pay for such safety is that it requires more DNA to be included in the re-integration construct. All the deleted upstream sequences must be included in the re-integration vector to ensure native expression (see next section).

For many genes, the translation initiation codon is not located in the first exon but further downstream. In such cases, it is a good idea to choose a 5’ breakpoint in the intron preceding the initiation codon, as this facilitates the subsequent creation of conditional alleles and reporters. As with the 5’UTR, it is generally safer to keep some distance between the intronic attP site and the downstream splice acceptor (~100bp). And, as before, it is advisable to choose insertion sites that are poorly conserved with other *Drosophila* species. To ensure proper splicing and expression, it is important that the sequence located between the insertion site and the translation initiation site be reconstituted in the rescuing construct. This approach, which we used to modify the *dpp* gene [15] is also described in the subsequent section.

In general, it is preferable to delete whole coding exons (as opposed to introducing breakpoints within coding sequence). This is because subsequent design of conditional alleles requires the insertion of FRT sites, which could interfere with protein activity if present in coding sequence. Remember however that, as discussed above, deletion of whole exons could allow alternative in-frame translation initiation sites located in downstream exons to be used.

### Insertion of various genetic elements in ΔattP alleles

Insertion-ready deletion (ΔattP) alleles are invaluable because they enable genetic elements to be expressed in the same pattern as that of the endogenous gene. In the previous section, we have described examples of re-integration that highlight the need to carefully select the deletion breakpoints. Below we provide an overview of how re-integration of various DNA fragments can contribute to insights about gene activity and function. We have previously described vectors for integration of DNA fragments into the attP site (RIVs) ([13]). With these vectors, successful integrants are selected with *Pax-Cherry* or *white*^+^ as a genetic marker. We have since generated RIV’s carrying *Pax-GFP* (Fig. 3D; Table 1) to allow selection of integrants into an attP site generated with a targeting vector marked with *Pax-Cherry*. The availability of distinct genetic markers for initial gene targeting and re-integration obviates the need to remove the first marker before proceeding to re-integration. Both markers can then be removed at once with one Cre-mediated step, thus saving time.

**Fig. 3.**
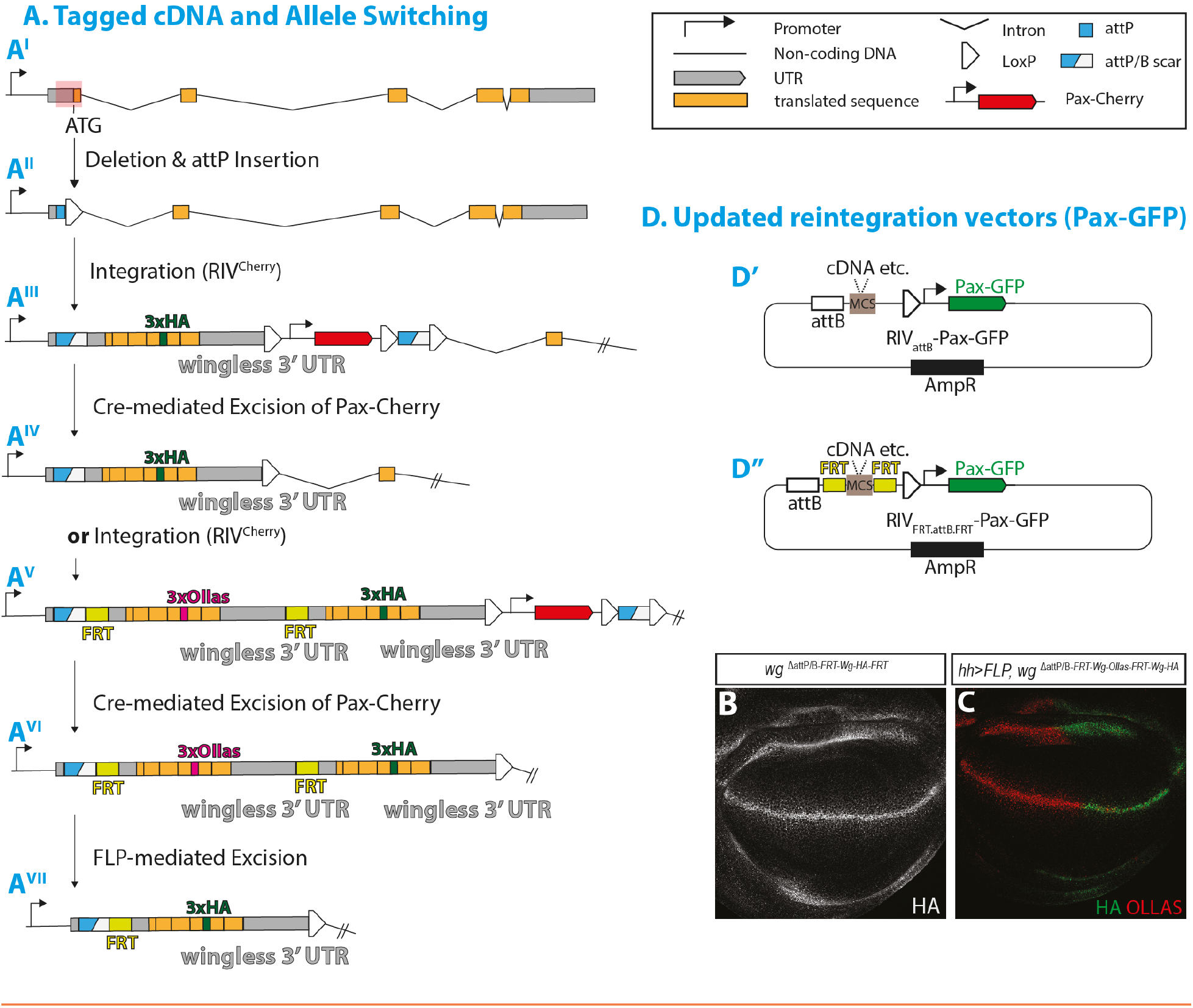
Insertion of various genetic elements in ΔattP alleles. (A) The *wingless* locus after replacement of the first coding exon by an attP site and removal of the *Pax-Cherry* cassette (A^I^ and A^II^, as shown in Fig. 2D). Locus configuration after re-integration of HA-tagged *wingless* cDNA (incl. wingless 3’UTR, A^III^). The *Pax-Cherry* cassette can then be excised by crossing to a Cre-expressing strain (A^IV^), allowing HA-Wingless to be expressed in a seemingly normal fashion as indicated in panel B. Also shown is an allele carrying an FRT-excisable *Ollas-wingless* cDNA (+ *wingless* 3’UTR) followed by *HA-wingless* cDNA (+ *wingless* 3’UTR) (A^V^). As with the *HA-wingless* allele, the *Pax-Cherry* cassette used for integration into the attP site can be removed by Cre (A^VI^). In the resulting strain, the gene product switches from Ollas-tagged to HA-tagged Wingless following excision of the FRT cassette (A^VII^, allele switching, see panel C). Here we used our previously published RIV^Cherry^ (with Pax-Cherry cassette) or RIV^FRT MCS FRT MCS3^ (also marked with *Pax-Cherry)* for re-integration ([13]). (B) Wing imaginal disc from a *wg*^ΔattP/*B-FRT-Wg-HA-FRT*^ larva (as described in A). (C) Wing disc from a *wg*^ΔattP/*B-FRT-Wg-Ollas-FRT-Wg-HA*^ larva after *hedgehog-Gal4/UAS-FLP* mediated allele switching (as described in A). (D’) Generic re-integration vector with *Pax-GFP* as a genetic marker. This was designed to allow integration in the attP site introduced in the genome with TV_ΔattP_-Pax-Cherry vector without the need for prior floxing out *Pax-Cherry*. Both markers can be floxed out simultaneously after re-integration. (D”) *Pax-GFP* marked re-integration vector with FRTs flanking a multiple cloning site (MCS) – this vector allows the creation of conditional alleles.

As mentioned above, re-integration of the deleted DNA can be used to validate the original mutation. In some cases, rescue can be achieved with a cDNA, if a small deletion (leaving regulatory introns in place) suffices to abrogate gene activity. Whether rescue is achieved with a cDNA or genomic DNA, DNA encoding a tag can be inserted in the rescue construct so that the protein product can be tracked at endogenous level. This is illustrated for HA-Wingless ([16], Fig. 3A, B). One important advantage of insertion-ready deletions is that they enable the creation of conditional alleles by flanking the rescue fragment with FRT sites. Such alleles are particularly useful to abrogate gene activity in specific cells/tissues at experimentally defined times. Of course, it is also possible to create tagged conditional alleles, as we have done before for *Dpp* ([15]). In another form of conditional allele, DNA encoding a modified or tagged form of the protein becomes expressed upon deletion of the FRT-flanked rescuing fragment, typically expressing the wild type form ([14, 16]). This approach is illustrated in Fig. 3A, C with the locus switching from expressing Ollas-tagged to HA-tagged endogenous Wingless protein. Note that in this case, the native 3’UTR must be appended to both coding sequences to ensure proper expression. This allele-switching approach would be particularly useful to assess the phenotype of mutant proteins that are expected to be dominant lethal (such as for example a phospho-mimetic).

Insertion-ready deletion alleles also allow the creation of reporters similar to those described in the section on insertion-only alleles, albeit in a somewhat more laborious way (since two steps are required and additional sequences might be needed to ensure expression). One example of a transcription reporter created by integration of GFP-encoding DNA into the attP site is a previously described *hid-GFP* ([17]). We also inserted other reporters (including tagRFP, LexA, and QF2) in *hid, reaper* and *grim* (stocks available upon request). Thus, any coding sequence can be expressed under the control of the targeted gene. Remember however that, to ensure proper expression, upstream genomic sequences (e.g. parts of the 5’UTR or an intron) that were deleted by the original mutation must be included in the reintegrated DNA. This can be achieved by appending these sequences in the 5’ primer used to amplify these fragments for insertion into the basic re-integration vectors, e.g. RIV_attB_-Pax-GFP (Table 1, Fig. 3D’) or RIV^white^ and RIV^cherry^ [13]. Alternatively, to express Gal4 from the locus, the deleted sequence can be inserted upstream of the Gal4 coding region of RIV^Gal4^ ([13]).

### Mistaken confirmation of the intended deletion

Phenotypic rescue by re-integration is the gold standard for verification that a given mutation causes a phenotype. However, as we found, such an assay can be misleading. In one project, we aimed to delete *Rpl41*, which encodes a component of the 60S subunit of the ribosome. The deletion was confirmed by PCR (data not shown) and, in the resulting homozygous larvae, highly proliferating tissues such as the brain and imaginal discs were abnormally small, a phenotype expected from ribosomal dysfunction (Fig. 4A-E). The deleted coding sequence was then reintroduced via the attP site and this apparently rescued the phenotype (data not shown). However, further analysis showed that the original phenotype was due to a mutation in *APC5*, which encodes a component of the anaphase promoting complex. It appears that the chromosome we initially created harbored a mutation in both *Rpl41* and *APC5* and that the *APC5* mutation segregated away from the *Rpl41* allele during the crosses that followed re-insertion of the *Rpl41* cDNA. We cannot determine whether, in this example, the *APC5* mutation was the result of an off-target effect or pre-existed in the host strain. Nevertheless, the lesson of this episode is that, for previously uncharacterized genes, it is necessary to verify that the phenotype segregates with the *Pax-Cherry* marker.

**Fig. 4.**
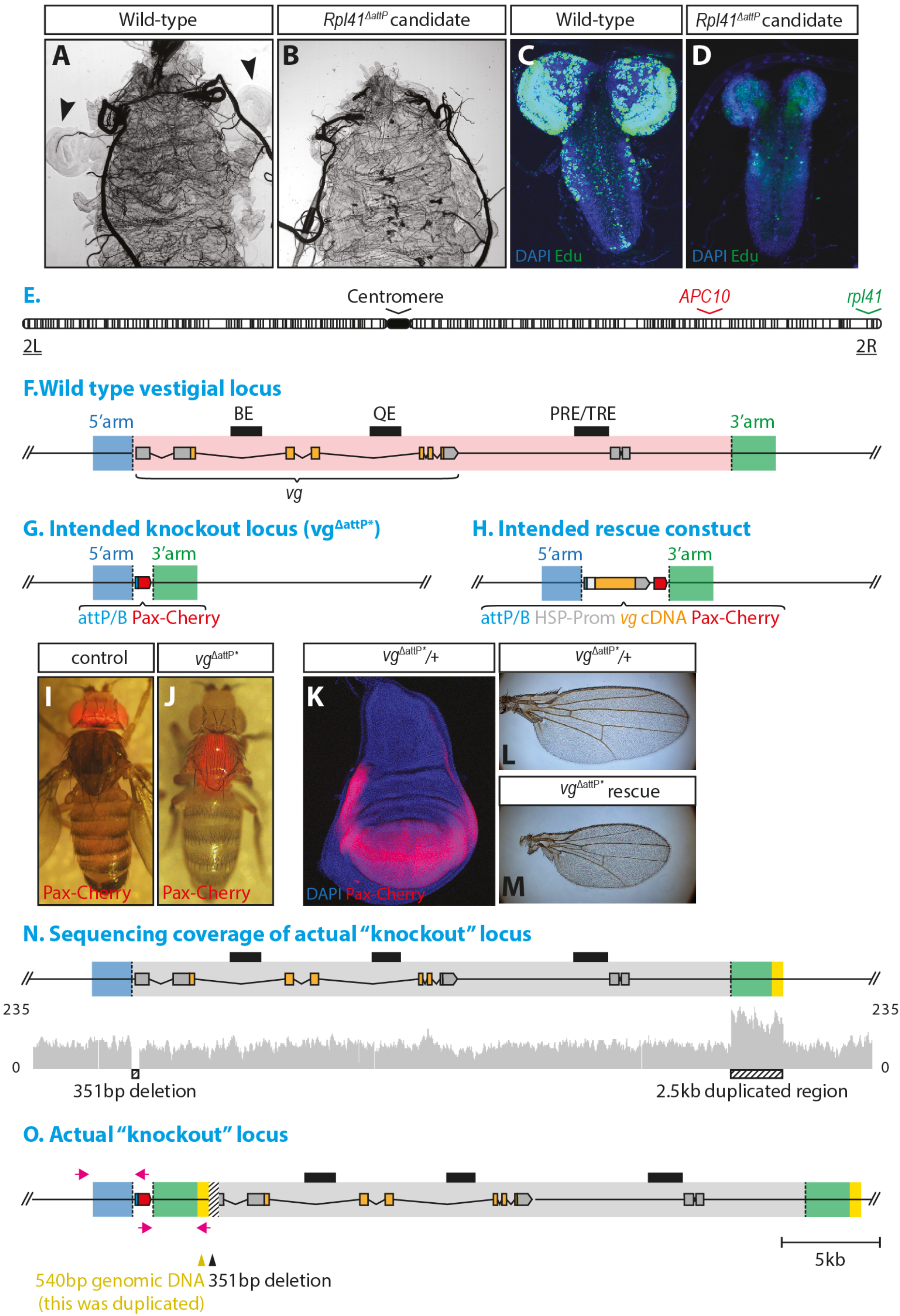
Mistaken confirmation of the intended deletion. (A, B) Wild type and *Rpl41*^ΔattP^ candidate mutant larva at third instar. Larvae have been turned inside-out to expose wing discs (arrowheads). Note the small size of wing discs in the *Rpl41*^ΔattP^ candidate mutant larva. (C, D) Brain of a *Rpl41*^ΔattP^ candidate mutant and wild type at third instar. Note the small size and reduced proliferation in the mutant tissue, particularly in the optic lobes. (E) Heterochromatin map of chromosome 2 showing the positions of *rpl41* and *APC5* loci. (F) The *vestigial* locus with the coding exons in orange, untranslated sequences in grey and the intended deletion in red box. The 5’ arm and 3’ arm used for homologous recombination are indicated in blue and green respectively. BE, QE and PRE/TRE are previously characterised regulatory elements [22] [23]. (G) Expected organisation of the *vestigial* locus following homologous recombination with the homology arms shown in F. This allele will be referred to as *vg*^ΔattP*^, with the * indicating that the actual deletion is not as expected. (H) Expected organisation of the *vestigial* locus after re-integration of the *vestigial* cDNA at the attP site (with *Pax-Cherry* as the visible marker of re-integration). (I) Expression of Cherry in a Pax-Cherry control fly, indicating the normal pattern of Pax-Cherry expression. (J) Cherry expression in *vg*^ΔattP*^ fly. (K) Cherry expression in a *vg*^ΔattP*^/+ wing imaginal disc. (L, M) Re-integration of *vestigial* cDNA into *vg*^ΔattP^* fully rescued gene function as evidenced by the wild-type appearance of wings. (N, O) Whole genome sequencing revealed that the 27 kb region that we thought to be deleted was merely displaced by about 3kb. This rearrangement was accompanied by a 351 bp deletion, which disrupted the expression of the native *vestigial*, but left most regulatory sequences intact. PCR primers that produced mis-leading PCR products are shown in magenta.

Another instance of mistaken confirmation of a CRISPR-induced deletion provides a useful lesson. Here we aimed to delete the whole *vestigial* locus, which spans 27 Kb of genomic DNA (gene editing strategy outlined in Fig. 4F-H). The resulting flies lacked wings, as expected from *vestigial* loss of function and PCR amplification with primers spanning the homology arms suggested that the intended modification was achieved (Fig. 4J, data not shown). Surprisingly however, the *Pax-Cherry* marker was found to be expressed in the same pattern as that of *vestigial* (Fig. 4I-K). We therefore considered the possibility that a re-integrated *vestigial* cDNA would be similarly expressed and thus possibly rescue the phenotype. Indeed, flies resulting from such re-integration had smaller but properly patterned wings despite the presumed absence of all known regulatory elements (Fig. 4L, M). In light of this surprising result, we considered the possibility that the deleted DNA might have translocated elsewhere in the genome and proceeded with whole genome sequencing of the original deletion strain. This showed that the 27 kb region was not deleted as originally thought but instead was displaced by about 3kb, with partial duplication of the 3’arm (Fig. 4N, O). We suggest that large deletions could be prone to complex rearrangements and that such modifications must be confirmed with multiple independent PCR assays (covering the putative deletion).

### Engineering regulatory regions

So far, we have only described procedures that modify the coding region. We have also gained some experience in creating designer alleles that modify the regulatory region. One lesson from this experience is worth sharing. Our aim was to assess the relevance of two AP-1 and two Schnurri binding sites located in the regulatory region of *reaper* (Fig. 5). The AP-1 sites mediate transcriptional activation by JNK signaling while the Schnurri sites mediate repression by Dpp signaling. We found that deletion of the four sites (Δ_4sites_=*reaper*^Δ*attPΔ1.9kb*^, Fig. 5C) was dominant lethal, as suggested by the finding that all (n=17) recovered F1 heterozygous animals (recognized by expression of the white+ marker) died at pupal stage with apparent massive degeneration of internal organs. Tellingly, a larger deletion encompassing the same regulatory region and the coding sequence was heterozygous viable (though homozygous lethal). This suggests that the dominant lethality of the Δ_4sites_ mutation is caused by ectopic expression of *reaper* following the deletion of essential repressive elements (probably the Schnurri sites). Therefore, removal of repressive elements is likely to cause dominant mutations that are difficult to recover. Assessing their function might therefore require a conditional approach by flanking them with FRT sites.

**Fig. 5.**
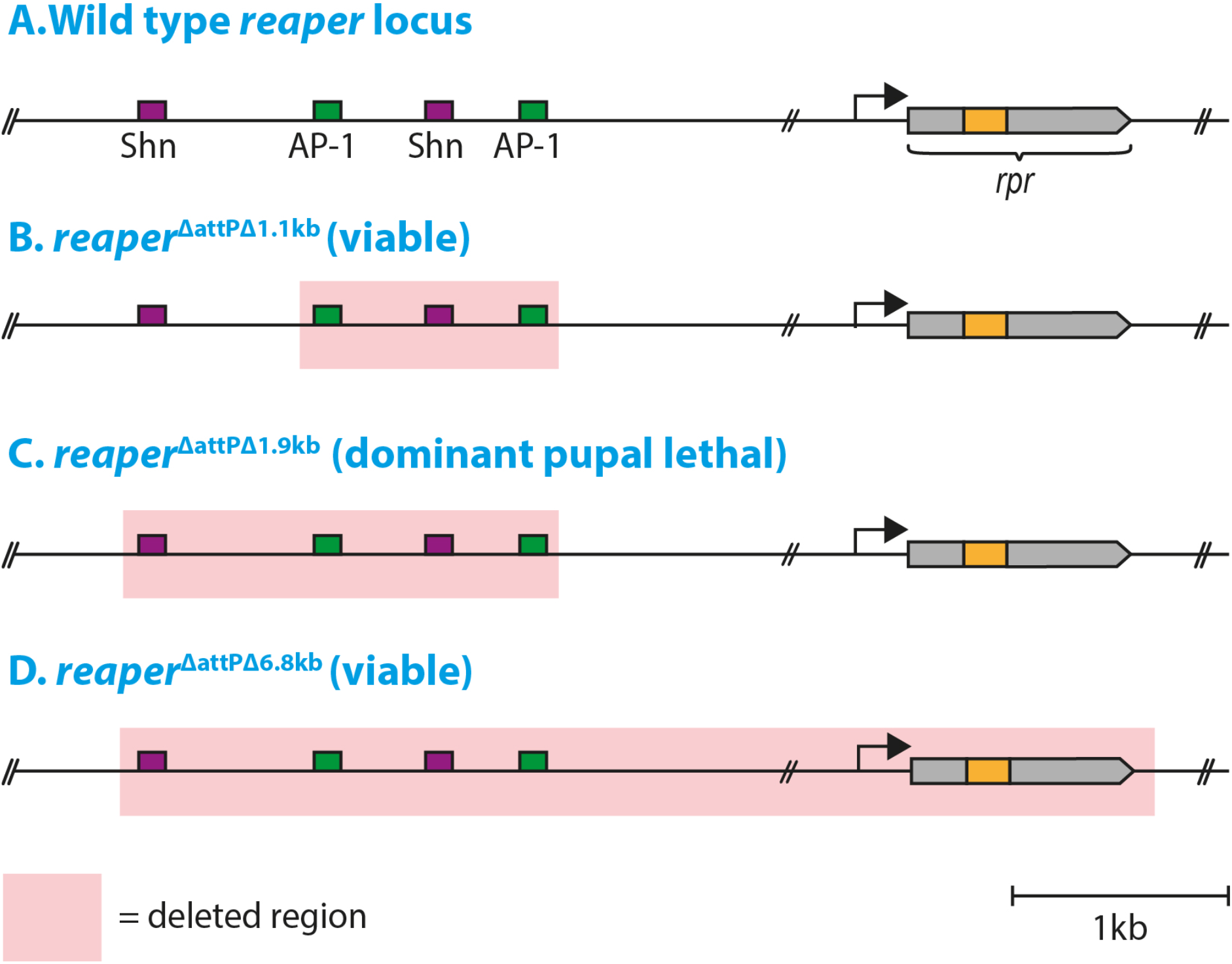
Engineering regulatory regions. (A-D) The *reaper* locus, with the various deletions shown in red shading. Known regulatory elements are marked in purple (Schnurri binding sites, Shn, which respond to Dpp) and green (AP-1 binding sites, which mediate the response to JNK).

## Conclusion

Thanks to CRISPR/Cas technology, most genes have become accessible to reverse genetics, especially in model organisms. However, obtaining a loss-of-function allele is only the first step towards understanding gene function. More sophisticated tools to measure gene expression and to control gene activity at experimentally defined time and space are needed. We have shown how other means of DNA manipulations (site-specific recombination and integration) can be combined with CRISPR/Cas to generate such tools. As we found, many factors must be considered to ensure that the achieved genomic modification serves the intended purpose. We highlight instances when gene inactivation is incomplete or integrated DNA fragments are not expressed. Heeding these lessons should enable the creation of genetic tools for an unprecedented range of experimental approaches.

## MATERIALS AND METHODS

All newly described plasmids were constructed by standard molecular techniques, including Gibson assembly ([24]). They will be deposited at Addgene.

Flies were reared at a consistent density on standard cornmeal/agar media at 25°C, Wing imaginal discs were fixed in 4% PFA for 20 minutes and washed in PBS. Immunofluorescence staining was preceded by permeabilization in 0.3% PBT, and blocking in 2% NDS. Antibodies used in this study were mouse anti-Wingless (1:500; 4D4 DSHB), rabbit anti-HA (1:500; Cell signaling) and rat anti-Ollas (1:10; Novus Biologicals). The protocol for Edu incorporation is described in [25]. Embryonic cuticles were prepared as in Alexandre et al. [26]. In situ hybridization for *wingless* mRNA was carried out according to standard protocols [27].

## ACKNOWLEDGEMENT

We thank Yohanns Bellaiche for discussion and the plasmids of IO_@N_TV-Pax-Cherry and IO_@C_TV-Pax-Cherry (929 N- and C-ter linkPaxRed).

## AUTHOR CONTRIBUTIONS

CA designed and built all the plasmids with initial input from L B-L. IP, SC, JK, and HN generated *Drosophila* strains. The figures were assembled by IP, SC and CA. The manuscript was written by JPV with help from IP and CA.

